# Investigating the therapeutic potential of soluble TREM2 interactions with transgelin-2 in Alzheimer’s disease

**DOI:** 10.64898/2026.10.30.763206

**Authors:** David Saeb, Emma E. Lietzke, Evan A. Wood, Ana F. Costa, Emma C. Aldrich, Daisy I. Fuchs, Ashkon E. Chaghajerdi, Kimberley D. Bruce, Kayla G. Sprenger

## Abstract

Triggering receptor expressed on myeloid cells 2 (TREM2) is a regulator of microglial function in Alzheimer’s disease (AD), and its soluble ectodomain (sTREM2) has emerged as a potential neuroprotective factor. Recent work shows that sTREM2 exerts protective effects through direct interaction with neuronal transgelin-2 (TG2) and modulates downstream tau pathology, yet the molecular basis of this interaction remains unknown. Here, we use all-atom molecular dynamics simulations to define the structural mechanisms of sTREM2–TG2 binding. We identify an electrostatically driven interface centered on the TG2 calponin homology domain. This surface is engaged by a binding motif on sTREM2 in which charged anchoring residues are flanked by hydrophobic contacts. We further show that the AD-risk TREM2^R47H^ variant maintains a more open, solvent-exposed CDR2 loop that redistributes interfacial contacts while preserving overall binding affinity, offering a structural explanation for reports that R47H does not exacerbate tau pathology in tauopathy models. Leveraging these insights, we design optimized sTREM2-derived peptides that outperform a previously reported therapeutic peptide in binding persistence, energetics, and recapitulation of native TG2 contacts, supporting engagement at the epitope through which sTREM2 suppresses tau phosphorylation. Taken together, our results define a mechanistic framework for the sTREM2–TG2 axis in AD and establish a structure-guided strategy for developing peptide modulators of this interaction.

## Introduction

Alzheimer’s disease (AD), the leading cause of dementia, affects millions worldwide and is projected to reach pandemic proportions, impacting over 150 million people by 2050^1^. AD is a progressive and fatal neurodegenerative disorder characterized by memory loss, cognitive decline, and impaired executive function, posing a major challenge to global healthcare systems. While the amyloid cascade hypothesis—centered on the accumulation of insoluble, extracellular amyloid beta (Aβ) plaques as a source of neurotoxicity in AD^2^—has long dominated AD research and achieved recent clinical successes in targeting Aβ^3–5^, it is evident that AD is a multifaceted disease. Beyond Aβ plaques, AD pathology involves intracellular neurofibrillary tangles (NFTs) associated with tau hyperphosphorylation and chronic brain inflammation driven by overactive microglia, the resident immune cells of the central nervous system^6,7^. Microglia play a dual role in AD, both mitigating and exacerbating pathology^8–11^. Protein aggregates and neuronal stress signals trigger microglial activation, leading to inflammatory responses^9,11^. In early AD, microglia can effectively clear Aβ plaques; however, chronic inflammation creates a positive feedback loop of neuronal damage and death, accelerating disease progression^12,13^. Elucidating the complex interplay among these pathological hallmarks is crucial for advancing AD research and therapeutic development.

A key mediator of neuroinflammation is Triggering receptor expressed on myeloid cells 2 (TREM2), a 230 amino acid (aa) transmembrane protein found on the surface of myeloid cells, with microglia as its sole site of expression in the central nervous system^14^. Structurally, TREM2 consists of an extracellular immunoglobulin (Ig)-like ligand-binding domain (19–130 aa), an unstructured extracellular stalk (131–174 aa), a single-pass transmembrane helix (175–195 aa), and an intracellular cytosolic tail (196–230 aa)^15^. In AD, TREM2 expression is upregulated, facilitating microglial recruitment to Aβ plaques^16–19^. Misfolded Aβ oligomers, which are precursors to plaques, are high-affinity ligands of TREM2. Oligomer-TREM2 binding triggers TREM2-mediated signaling^20,21^, which is critical for microglial fitness and function^22–24^, ^8–20^ Furthermore, this signaling facilitates a TREM2-dependent activation towards an AD-specific disease-associated microglia (DAM) phenotype that may be protective in early stages by enhancing plaque containment and clearance, but detrimental when chronically sustained^13,25–27^. Furthermore, TREM2 variants with loss-of-function mutations, particularly the R47H variant, significantly increase AD risk^28–30^ by likely impairing these early protective responses^31–34^. TREM2’s involvement in AD pathology extends beyond its interactions with Aβ, as it has also been implicated in tau phosphorylation. However, its contribution in this context remains unclear due to conflicting evidence supporting both exacerbating and ameliorating effects^35^. These discrepancies underscore the need to further elucidate how TREM2 interfaces with multiple AD hallmarks.

This complexity is further compounded by the existence of TREM2’s soluble form (sTREM2), which functions as a ligand and is mechanistically distinct from its canonical signaling activity as a membrane-bound receptor. As such, the Ig-like domain of TREM2 can be cleaved between His157 and Ser158 by ‘a disintegrin and metalloprotease’ (ADAM) family sheddases, releasing soluble TREM2 (sTREM2) into the extracellular space^36,37^. Additionally, sTREM2 can be generated through alternative splicing of TREM2 pre-mRNA^38^. The role of sTREM2 in AD pathogenesis is an area of active investigation. Current evidence suggests that sTREM2 exerts neuroprotective effects in AD^39,40^, including facilitating plaque clearance by binding Aβ plaques^41–43^, modulating neuroinflammatory responses^41,44^, and efficiently inhibiting Aβ fibril formation^45^. Beyond its potential neuroprotective roles, sTREM2 has emerged as a promising biomarker for AD^46^, with elevated cerebrospinal fluid levels reported in symptomatic disease relative to healthy individuals and changing dynamically across the disease continuum in association with tau pathology and neurodegeneration. Notably, higher sTREM2 levels correlate with slower disease progression, reflected in reduced symptom presence and brain atrophy^47^.

Yet, these levels also fluctuate during disease progression, with initial declines during asymptomatic Aβ accumulation, increases during tau aggregation, and reductions during dementia progression^48–50^. Such dynamics highlight sTREM2’s complex role in AD pathogenesis and its potential as both a biomarker and therapeutic target.

Beyond sTREM2’s emerging therapeutic relevance, recent work suggests that sTREM2 may also act directly at the neuronal surface. Zhang et al. revealed that sTREM2 may exert neuroprotective functions through direct interaction with transgelin-2 (TG2; not to be confused with transglutaminase 2), a versatile protein expressed in neurons and other diverse cell types, including smooth muscle and immune cells^51^. Although TG2 is primarily cytoplasmic and lacks a canonical transmembrane domain, it is thought to associate with membranes under certain conditions, where it can interact with extracellular ligands and enable receptor-like activity via its calponin homology (CH) domain^51–53^. Zhang et al. demonstrated that sTREM2 directly interacts with membrane-associated TG2, leading to downstream inhibition of the neuronal RhoA–ROCK pathway. This inhibition reduced tau phosphorylation, a key driver of tau fibrillization and neuronal toxicity. Consequently, tau aggregation decreased, mitigating neurocognitive effects in a tauopathy mouse model^51^. These findings suggest that the interplay between sTREM2 and TG2 holds promise for developing targeted AD therapies that are unique from those targeting membrane-bound TREM2. However, the precise molecular mechanisms underlying TG2-sTREM2 interactions remain incompletely understood, and the rational development of targeted therapies will require deeper mechanistic insight.

Building on these findings, we aimed to comprehensively characterize the molecular-level interaction between sTREM2 and TG2. Through fully atomistic molecular dynamics (MD) simulations, we investigated the key binding surfaces and residues of sTREM2 and TG2 mediating this interaction. By employing long-timescale MD simulations, we sought to capture the behavior of the sTREM2–TG2 complex over biologically relevant timescales, revealing the electrostatic and hydrophobic determinants of binding and how the R47H variant reorganizes this interface through CDR2 loop dynamics. Additionally, we simulated a therapeutic peptide sequence proposed by Zhang et al. and optimized a novel peptide to enhance its efficacy in targeting this interaction. Ultimately, our work aims to deepen understanding of AD pathogenesis and identify new therapeutic strategies.

## Methods

### Preparation of Simulated Structures

To study wildtype (WT) sTREM2, we utilized the AlphaFold-predicted structure (AF-Q9NZC2-F1) of full-length TREM2^54,55^, simulating residues 19–157. This structure was selected for its inclusion of the flexible stalk region and strong alignment of the Ig-like domain with experimentally derived structures. The R47H point mutation was modeled by combining an experimentally resolved crystal structure of the Ig-like domain (Protein Data Bank (PDB) 5UD8) with the unstructured stalk from the AlphaFold-predicted structure^56^. Previously, we conducted all-atom MD simulations of these structures and clustered their trajectories to generate initial docking conformations using the Gromos method implemented in GROMACS.

For TG2, we used the only available experimentally derived structure (PDB 1WYM), which includes the CH domain and partial unstructured N-and C-termini^57^. We simulated this structure without incorporating the remaining unstructured termini, as it fully encompasses the primary binding epitope. TG2 therapeutic peptide candidates were isolated as segments of the AlphaFold-predicted TREM2 structure. Three therapeutic peptide sequences, designated Peptide1, Peptide5, and Peptide6, were identified based on Zhang et al.’s findings and the results of WT sTREM2-TG2 simulations, as detailed in the Results section^51^.

### Molecular Dynamics Simulations

In total, four pure-component systems (TG2 and three therapeutic peptides) were simulated prior to docking studies for 1 μs each. Simulations of WT and R47H sTREM2 were described in a previous study. All systems, including proteins and counterions, were parameterized using the CHARMM36^58,59^ forcefield implemented in GROMACS^60^. The TIP3P water model was used to simulate explicit solvent^61^. All systems underwent steepest-descent energy minimization, first in vacuum and then in solvent. Equilibration was performed with a 1 ns NVT simulation at 310 K using the Bussi-Donadio-Parrinello thermostat^62^, followed by a 1 ns NPT equilibration at 310 K and 1 bar using the same thermostat and Berendsen barostat^63^. Production simulations were conducted at 310 K and 1 bar, employing the same thermostat and Parrinello-Rahman barostat^64^, with a duration of 1 μs for each pure-component system.

Simulations employed the LINCS algorithm to constrain bonds involving hydrogen atoms^65^, with long-range electrostatics calculated using particle-mesh Ewald (PME) summations and a cutoff of 1.2 nm^66^. Lennard-Jones interactions were evaluated up to 1.2 nm and shifted to eliminate energy discontinuities, while neighbor lists were updated every 10 steps with a 1.4 nm cutoff. All simulations used a 2 fs timestep, with periodic boundary conditions applied in all directions. Combined systems (TG2 with sTREM2/peptides) used identical parameters during the 150-ns production simulations. These simulations were initialized using outputs from the pure-component trajectories, which were combined through molecular docking (see below).

### Molecular Docking Calculations

Prior to docking, the 1-µs trajectory of each pure-component system was clustered using the Gromos method implemented in GROMACS to obtain representative structures across the entire trajectory. Docking of TG2 to sTREM2 structures and therapeutic peptide candidates was performed using the LZerD protein-protein docking webserver^67–69^. LZerD employs geometric hashing with 3D Zernike descriptors to generate *de novo* structures and has been rigorously validated through the CAPRI performance evaluation^70,71^. In the docking calculations, TG2 was designated as the receptor, as it is membrane-associated, while sTREM2 and therapeutic peptides were treated as ligands. For each protein-protein system, docking was conducted with 2-3 clustered structures for each component. All clusters were cross-docked with one another to ensure that all relevant conformations were sampled. Docked complexes were analyzed based on the LZerD score, the number of conformationally similar complexes, and biological relevance. Using this approach, nearly all unique docked complexes were selected for subsequent MD simulations. This resulted in 9 WT sTREM2–TG2 models, 10 R47H sTREM2–TG2 models, 9 Peptide 1–TG2 models, 9 Peptide 5–TG2 models, and 8 Peptide 6–TG2 models for further MD simulations.

### Trajectory Analysis Protocols

Simulation convergence was assessed using root-mean-square deviation (RMSD) calculations for both the complexes and individual proteins. To calculate binding free energies, we used the Molecular Mechanics Poisson-Boltzmann Surface Area (MM-PBSA) method implemented in gmx_mmpbsa^72,73^. MM-PBSA calculations were performed over the last 50 ns of the trajectories. Fractional occupancies of sTREM2 onto TG2, TG2 onto sTREM2, and each therapeutic peptide onto TG2 were computed by calculating the fraction of trajectory frames in which a given component was within 4 Å of another throughout the simulation. Due to the highly dynamic nature of peptide binding and frequent binding/unbinding events, our Peptide-TG2 simulations occasionally captured very short transient self-interactions across periodic boundaries during the first 100 ns. To avoid artifacts arising from these events, all fractional occupancy analyses were therefore restricted to the final 50 ns of each Peptide–TG2 trajectory. Protein conformational shifts and stability were evaluated via RMSD and root-mean-square fluctuation (RMSF) analyses, while visual analysis of the simulation trajectories was performed using Visual Molecular Dynamics (VMD) and ChimeraX^74–77^. RMS(Δ) was calculated as the root mean square of the per-residue difference in fractional occupancy between each peptide–TG2 complex and the WT sTREM2– TG2 complex, computed over all aligned TG2 residues. Protein visualizations were rendered with ChimeraX^75–77^. MDAnalysis and MDTraj were used to perform further specialty analyses including DSSP and SASA^78–80^.

### Convergence Assessment of Full-Length sTREM2-TG2 Simulations

Because RMSD is referenced to the initial docked poses, lower RMSD values do not necessarily indicate more favorable binding poses but rather reduced deviation from the starting structure. Accordingly, we interpreted convergence as trajectories settling into bounded RMSD fluctuation windows, which, for full-length sTREM2-TG2 complexes, were generally reached early and persisted throughout the simulations, indicating broad convergence across models. Within these windows, fluctuations sometimes narrowed into more stable plateaus, but these were not directional—longer simulation times did not necessarily yield more favorable rearrangements, and in several cases the tightest RMSD plateaus occurred mid-trajectory rather than in the final frames. With this context, we evaluated the relative stability of full-length sTREM2-TG2 complexes by computing the percentage of simulation time each trajectory remained within a ±3 Å Cα RMSD window (calculated for the full protein complex while excluding TG2’s unstructured termini), assessed over 20 ns intervals. Across all WT simulations, Models 9, 8, 4, 5, 1, 7, and 3 exhibited convergence for over 50% of the time, listed here in descending order of convergence percentage, suggesting greater favorability for those initial docked conformations (**Supplementary Fig. 2A**).

Similarly, this analysis identified Models 9, 4, 3, 10, 2, 6, and 7 as the most stable R47H simulations (**Supplementary Fig. 3A**). This convergence-based stability metric was used to downsample the docking ensemble, retaining only the most stable models for relevant analyses.

## Results and Discussion

### Conserved binding domain of TG2 modulates direct interaction with WT sTREM2

Prior studies of TG2 have primarily focused on its role in asthmatic disease states^52,53,81^. We aimed to identify and compare how TG2 interactions with sTREM2 differ from those previously observed with TSG-12, a small-molecule TG2 agonist candidate identified in prior asthma studies, to elucidate TG2’s therapeutic role in AD. A 2023 study by Wu et al. conducted MD simulations on the full structure of TG2 complexed with TSG-12^81^. Similar to our approach, they utilized the crystal structure of the CH domain of TG2 (PDB 1WYM) but extended the partially resolved N- and C-termini by modeling in the missing terminal residues using homology modeling. Their findings demonstrated that the CH domain remained highly stable throughout the simulation, serving as the exclusive binding site for TSG-12. Because of this stability—and given that TG2’s C-terminus would likely be membrane-tethered in a physiological context—we excluded the unresolved terminal residues beyond those captured in the crystal structure, as these missing residues could artificially occlude the CH domain binding interface if modeled in. This also allowed us to focus exclusively on the conserved CH domain for our binding studies (**Fig. 1A**). Notably, Wu et al. identified 10 TG2 residues in frequent contact with TSG-12 (I31, P47, F52, R49, P45, E27, G48, Q46, R44, and Q28), of which three were found to be energetically essential for TSG-12 binding (E27, R49, and F52). Highlighting these residues on TG2 revealed a distinct Small Molecule Pocket (SMP) for TSG-12 binding (**Fig. 1A**).

**Figure 1.**
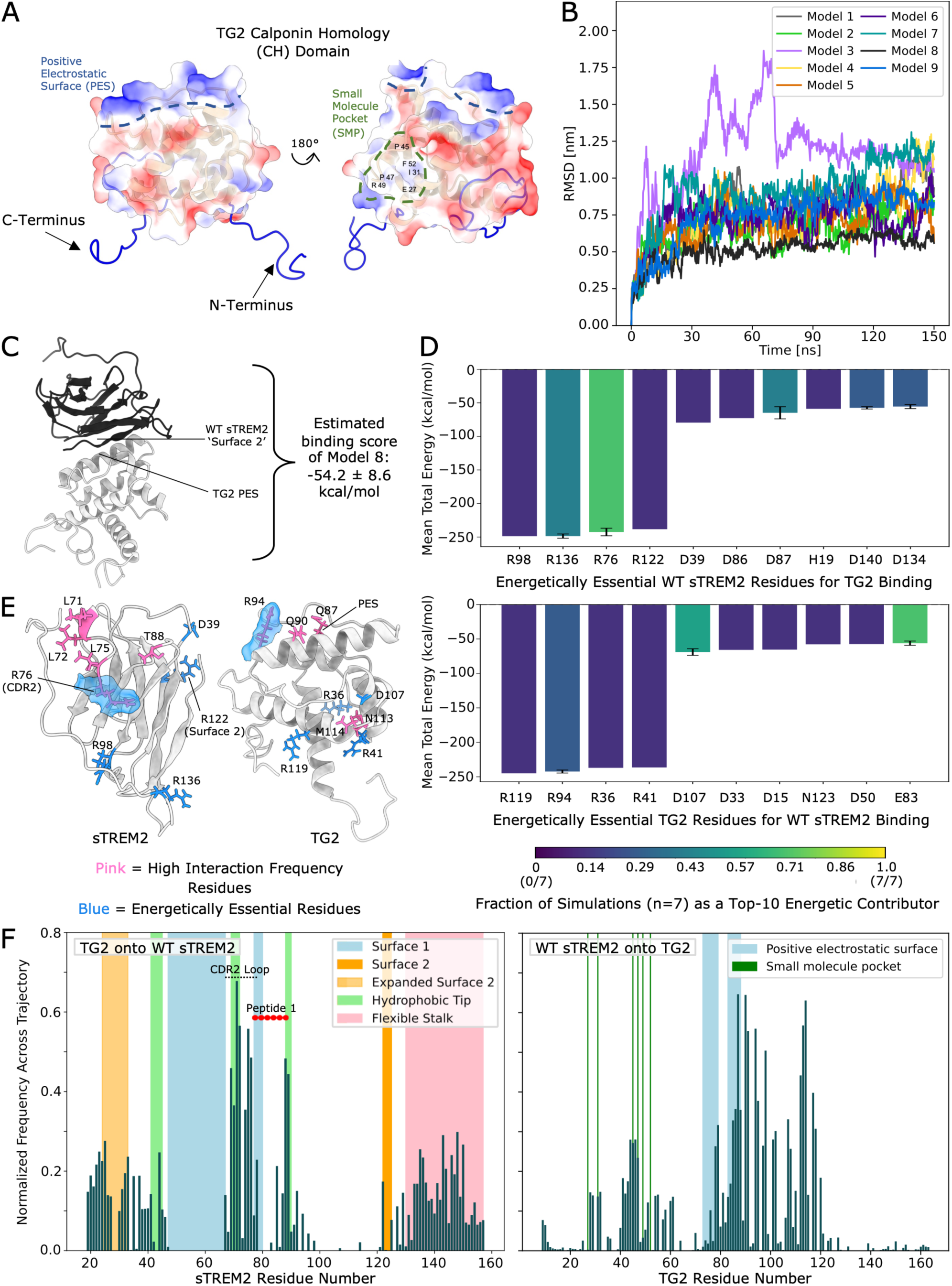
Structural and energetic basis of WT sTREM2–TG2 binding. (A) Electrostatic surface representation of TG2 highlighting a novel positive electrostatic surface (PES) and the previously characterized small-molecule pocket (SMP). The SMP is delineated by the top six TG2 residues most frequently contacting the small-molecule agonist TSG-12 in prior simulations (including the energetically essential residues E27, R49, and F52), as reported by Wu et al. Blue indicates positive electrostatic potential, red indicates negative electrostatic potential, and white indicates near-neutral regions. (B) Whole-system Cα-RMSD of WT sTREM2-TG2 complexes. (C) Visualization of the most energetically favorable WT sTREM2-TG2 complex (Model 8) with the corresponding MM-PBSA binding score and standard deviation, and relevant structural features labeled. (D) Residue-level MM-PBSA total energy decomposition for WT sTREM2 (top) and TG2 (bottom), aggregated across seven simulations with >50% convergence (see Methods). For each residue, only simulations in which the residue exhibits a strong stabilizing contribution (total energy ≤ −5 kcal/mol) are included in the calculation of the mean and standard deviation (error bars). Bar color indicates the fraction of simulations in which each residue meets this cutoff, reporting the consistency of its contribution across models. (E) sTREM2 (left) and TG2 (right) structures showing the top five residues with high interaction frequency (pink) and the top five residues with the strongest favorable energetic contributions from MM-PBSA analysis (blue). (F) Normalized fractional occupancy of WT sTREM2 residues by TG2 (left) and TG2 residues by WT sTREM2 (right) calculated across seven simulations with >50% convergence.

To elucidate the interaction between TG2 and sTREM2, we performed nine 150-ns MD simulations of unique WT sTREM2–TG2 complexes identified through a global molecular docking search with LZerD (see **Methods**). RMSD analysis indicated that seven of the systems converged rapidly, while the remaining two systems exhibited larger initial movements before stabilizing toward the end of the simulations (**Fig. 1B**, see Models 3 and 7 for all RMSD profiles, see **Supplementary Fig. 1A, C-E**). RMSF calculations revealed high per-residue fluctuations in sTREM2’s flexible stalk domain and TG2’s unstructured termini, indicating greater flexibility and dynamic motion in these regions, as expected (**Supplementary Fig. 1B, F-G**). In contrast, the Ig-like domain of sTREM2 and CH domain of TG2 both exhibited low RMSF values, which, together with low RMSD values, indicate high overall protein stability (**Supplementary Fig. 1F-G**). Importantly, the global stability of the Ig-like domain also extended locally to the second complementarity-determining region (CDR2) of sTREM2, which retained a closed-loop conformation akin to that in the starting crystal structure. This is consistent with our previously published results, which showed that the CDR2 loop is stabilized by the stalk and remains in this wconformation when bound to phospholipids (PLs)^42,82^. This behavior suggests that CDR2 structural stability is maintained across diverse binding partners.

We next assessed which conformations among our nine WT simulations were most conformationally stable and energetically favorable sTREM2–TG2 interactions (see **Methods** and **Supplementary Fig. 2A**). Visual trajectory analysis showed that sTREM2 and TG2 engaged through different interfaces across models (see **Supplementary Fig. 2B** for representative binding poses), each of which mostly remained locally stable during the simulations, without transitions to entirely new binding sites. However, interfacial repositioning at said binding sites occurred over the 150-ns timescale, reflecting distinct yet coexisting binding modes at a given interface. Among the models, Model 9 represented the most stable configuration, in which the CDR2 loop on sTREM2 engaged on and adjacent to a Positive Electrostatic Surface (PES) mapped to the ‘top’ of TG2 (**Supplementary Fig. 2A-B**). This convergence-based stability metric was then used to downsample the ensemble, retaining only the most stable models for downstream analysis.

Because our docking approach sampled broadly and multiple relatively stable sTREM2–TG2 complexes were identified during the simulations, we performed MM-PBSA binding free energy calculations (see **Methods**) to identify those yielding the most favorable binding affinities. The two overall most energetically favorable complexes (Models 8 and 3) both showed interactions between the PES on TG2 and the region of sTREM2 denoted Surface 2 in previous studies (**Fig. 1C**, **Supplementary Fig. 2B**). Given the high stability of these models (>50% convergence during simulations), these findings likely reflect the influence of the flexible stalk as a driving force for promoting alternative binding surfaces on sTREM2. This aligns with our past work that observed a shift in PL engagement from TREM2’s Surface 1 and CDR2 to Surface 2 in sTREM2. Despite the two most energetically favorable complexes (Models 8 and 3) engaging primarily through Surface 2, residue-level energy decomposition revealed that one of the strongest favorable energetic contributors on sTREM2 resided on CDR2 (R76), identified with high consistency across models, whereas the top-contributing residue on Surface 2 (R122) appeared in only Model 3 (**Fig. 1D**, top; **Fig. 1E**, left). Consistent with this, many of the most conformationally stable complexes (including the highest-convergence Model 9) featured direct CDR2–PES interactions (**Fig. S2B**). Taken together, these observations support prior work showing that both TREM2 and sTREM2 can diversely bind ligands due to variegated domains that are conformation- or ligand-specific (e.g., Aβ, PLs). Herein, Surface 2 offers a more favorable global binding interface due to reduced stalk competition, while CDR2 and hydrophobic tip residues consistently dominate local energetic contributions and are thus likely to be preferentially sampled and drive functional outcomes.

To more finely examine residue-level interactions beyond energetics and identify the molecular features governing their binding, we calculated average fractional occupancies of interacting residues in both TG2 and sTREM2 over the course of the simulations, averaging across the seven high-stability runs. Notably, CDR2 and hydrophobic tip residues exhibited higher interaction frequencies with TG2 residues than Surface 2 residues (**Fig. 1F**, left). While this may in part reflect that seven of our nine docked models had TG2 initially bound to CDR2 and adjacent regions (compared to two models with Surface 2), it also suggests that even at the docking stage, CDR2 and the hydrophobic tip presented a more accessible and favorable binding interface. This preference emerged and persisted even despite competition with the stalk for Surface 1 and CDR2 binding.

Notably, sTREM2 engaged TG2 in a pattern reminiscent of its interactions with PLs^42^. In addition to the CDR2 loop, sTREM2 interacted with TG2 via its hydrophobic tip, Expanded Surface 2 (newly defined in our recent study), and flexible stalk (**Fig. 1F**, left), suggesting that sTREM2 engages conserved regions across both lipid and protein binding partners. Strikingly, direct binding to the lower regions of Surface 1 was completely ablated, indicating that stalk-mediated occlusion may play an even larger role in sTREM2–TG2 engagement than in PL binding—potentially due to the larger size of TG2 or differences in electrostatic complementarity. This behavior contrasts with that of TREM2, which is known to engage ligands via Surface 1, and further reinforces that distinct surface engagement patterns govern sTREM2’s unique binding behavior. In sTREM2, the stalk actively alters and constrains surface accessibility through intermolecular interactions with the Ig-like domain, adding an additional layer of regulation absent in membrane-bound TREM2.

TG2’s interactions with sTREM2’s stalk further indicate that this region acts as a functionally significant structural element, enabling TG2 to distinguish between soluble and membrane-bound TREM2.

Fractional occupancy analysis of TG2 residues by sTREM2 revealed conserved binding to TG2’s PES and surrounding regions (**Fig. 1F**, right). Notably, sTREM2 had modest interaction with residues previously identified as mediating TSG-12 binding (small molecule pocket)^81^. Instead, sTREM2 primarily engaged residues on TG2’s opposite face, directly on and adjacent to the PES. Consistent with this, MM-PBSA calculations indicated that residues within and near the PES contributed most strongly to sTREM2–TG2 binding (**Fig. 1D**, bottom; **Fig. 1E**, right). This divergence from TSG-12’s binding pattern indicates that TG2 employs distinct surface sites and interaction chemistries for protein versus small-molecule binding partners, governed by complementary steric and electrostatic mechanisms.

Finally, we asked whether the residues most frequently engaging the binding partner overlapped with those making the strongest energetic contributions to binding. As such, we extracted the top five sTREM2 residues with the highest TG2 interaction frequencies from the fractional occupancy analysis, and vice versa. Initially, there appeared to be limited overlap between frequent residues and those contributing the strongest favorable energies. Both proteins’ energies were primarily derived from positively and negatively charged residues, emphasizing electrostatic interactions as key drivers of sTREM2–TG2 binding. Closer visual and sequence inspection, however, revealed that the high-occupancy residues tended to be less charged and/or more hydrophobic than the energetically essential residues, but often were situated directly adjacent to them. This arrangement suggests a local context or potential functional ‘motif’ in which charged residues provide strong energetic anchoring while flanking hydrophobic residues promote persistent contact, supporting efficient TG2 binding and stimulation by sTREM2. For example, R76 on CDR2 exemplifies this motif: it provides a strong favorable energetic contribution to sTREM2–TG2 binding and is positioned adjacent to hydrophobic, high-occupancy residues in the CDR2 loop (e.g., L71, L72, and L75).

Our analysis identified two regions with this motif: one on CDR2 and one spanning a broader domain across Surface 2 and Expanded Surface 2. This suggests that binding at both sTREM2 domains likely arises from shared biochemical features that enable both surfaces to engage TG2. Notably, residues R77–L89 comprising the sTREM2-derived therapeutic peptide proposed by Zhang et al. (Peptide 1; **Fig. 1F**, left, red circles) do not fully overlap with the sTREM2 regions showing the highest TG2 occupancies in our simulations (69–76 aa, 88–89 aa), although partial overlap provides confidence in the validity of our approach and results. Analysis of its sequence (RWNGSTAITDDTL) revealed a similar motif to that identified above, namely an arginine followed by a stretch enriched in hydrophobic residues, which may explain why the peptide retains TG2-binding activity despite only narrowly bridging the two peak TG2-occupancy regions. Together, these findings suggest that while Peptide 1 originates from a relevant sTREM2 region with the necessary biochemical motif and some high-occupancy residues, a rationally optimized peptide variant extending across one or both of the highest-occupancy regions could likely achieve even stronger TG2 binding. These findings establish a residue-level framework for sTREM2–TG2 engagement that we next apply to examine how the AD-risk R47H variant alters this interface, and to guide rational peptide design.

### The AD-risk mutation R47H enhances sTREM2–TG2 binding

Building on our characterization of TG2 interactions with WT sTREM2, we next asked whether disease-associated sTREM2 variants alter these binding dynamics and how this might inform the need for rationally optimized peptide design. We focused on the high-profile AD-risk mutation R47H to assess its impact on TG2 engagement. In previous work using amyloid precursor protein (APP) mouse models with seeded tau pathology, the R47H variant in human TREM2 was reported to reduce microgliosis around Aβ plaques while concomitantly increasing tau aggregation^83^. In contrast, in tauopathy-only PS19 mice expressing human TREM2, two independent studies reported that R47H did not exacerbate tau pathology: one found no significant effect when R47H was overexpressed in microglia^84^, whereas another reported attenuated tau phosphorylation in select brain regions, along with reduced brain atrophy and synapse loss in R47H versus WT TREM2^85^. These divergent outcomes are difficult to reconcile within a single mechanism, particularly because they have been attributed to opposing effects of R47H in microglia versus neurons. Because R47H lies in the Ig-like domain that becomes sTREM2, these models carry the variant in both the membrane-bound and shed forms, so microglial and soluble sTREM2 functions are perturbed together and cannot be disentangled. Given this, the emerging sTREM2–TG2 axis may be especially important, contributing to the possibly protective tau-related outcomes attributed to R47H via a novel neuronal-surface mechanism.

Given that the R47H mutation increased the flexibility of sTREM2’s CDR2 loop in our prior work^42^—reflecting destabilization of the loop’s open conformation by the stalk—we hypothesized that this increased flexibility may enhance sTREM2–TG2 engagement by maintaining an open, binding-competent CDR2 conformation, thereby modulating tau-related signaling in AD. To determine if this trend was preserved and perhaps accentuated upon TG2 binding, we performed ten 150-ns simulations of R47H sTREM2–TG2 complexes derived from our docking ensemble. Comparing the bound WT and R47H trajectories showed elevated RMSF values for both the stalk and CDR2 loop in R47H relative to WT sTREM2 (**Fig. 2A**), consistent with our previous unbound results.

**Figure 2.**
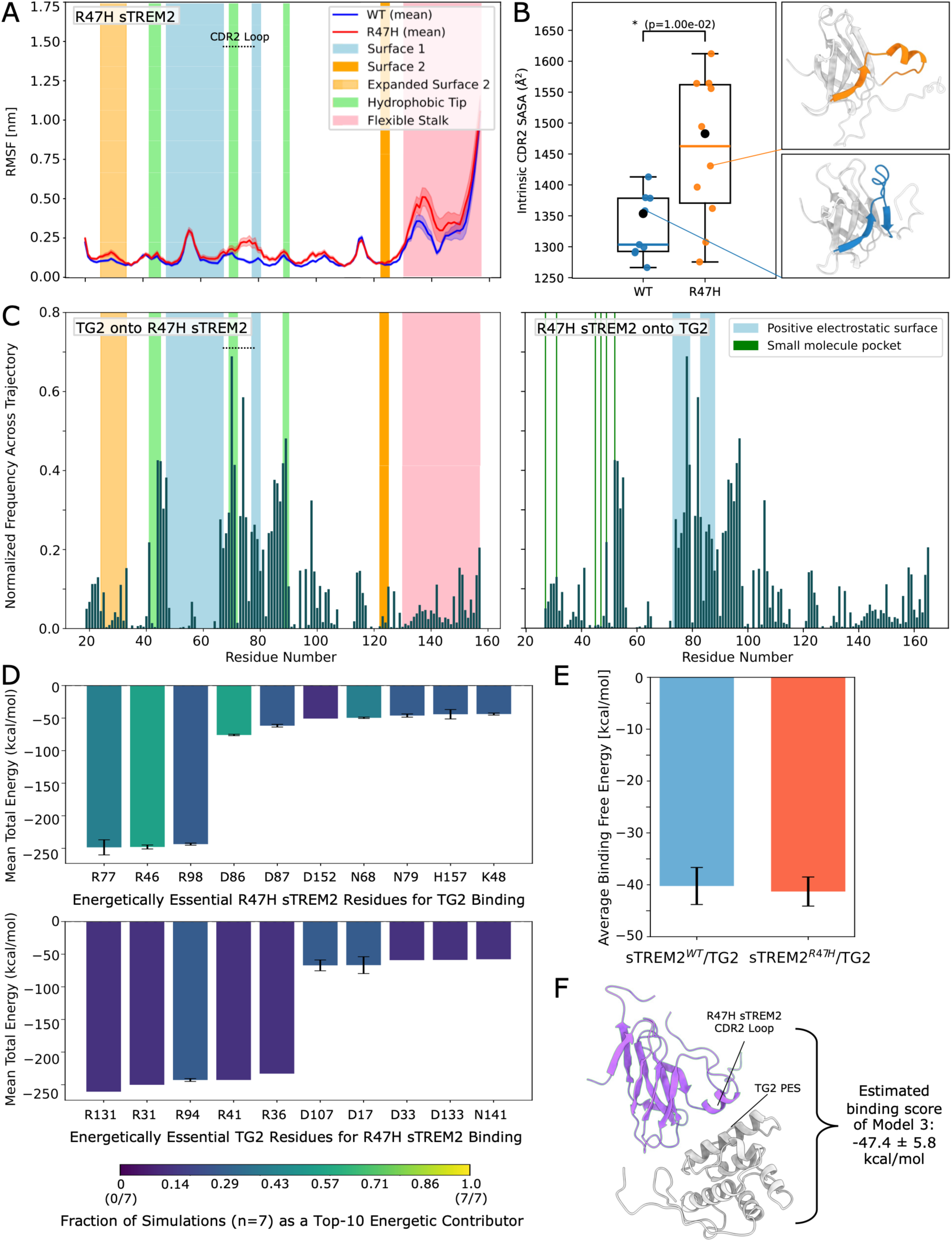
R47H-driven reorganization of the sTREM2–TG2 binding interface. (A) Cα RMSF profiles for sTREM2 WT (blue) and R47H (red) bound to TG2, shown as the mean ± the standard error of the mean (SEM) across all independent simulations. (B) Intrinsic solvent-accessible surface area (SASA) of the sTREM2 CDR2 region (residues 60–80) in WT and R47H models after removal of TG2. Boxplots summarize the distribution of per-replicate mean CDR2 SASA values, with individual per-replicate means shown as points; black points indicate values from previously published unbound sTREM2 simulations. Representative structures illustrating the closed (WT, blue) and open (R47H, orange) CDR2 conformations are shown on the right. Statistical significance was assessed using a Welch’s t-test (t = -3.03; p = 1.00e-2; n = 9 WT, 10 R47H). (C) Normalized fractional occupancy of R47H sTREM2 residues by TG2 (left) and TG2 residues by R47H sTREM2 (right) calculated across seven simulations with >50% convergence. (D) Residue-level MM-PBSA total energy decomposition for R47H sTREM2 (top) and TG2 (bottom), aggregated across seven simulations with >50% convergence (see Methods, Fig. 1). (E) Visualization of the most energetically favorable R47H sTREM2-TG2 complex (model 3) with the corresponding MM-PBSA binding score and standard deviation, and relevant structural features labeled. (F) MM-PBSA binding free energies for WT sTREM2–TG2 and R47H sTREM2–TG2 complexes averaged across seven simulations with >50% convergence. Bars show the mean and error bars indicate SEM.

Visual inspection confirmed that all bound R47H models maintained an open CDR2 loop throughout the simulations (**Supplementary Fig. 3C**), whereas bound WT models retained a closed conformation. To further assess whether this maintenance of a flexible, open CDR2 loop in the presence of R47H translated into a more exposed interface for TG2 binding, we calculated the solvent accessible surface area (SASA) of the broader CDR2 region (60–80 aa). This was calculated in all sTREM2 models after removing TG2 from the complexes, and in unbound simulations from our previous work as a control with no ligand present. Indeed, R47H models exhibited substantially higher CDR2 SASA than WT models on average (**Fig. 2B**, orange and blue dots, resp.). Moreover, when compared to our prior published unbound sTREM2 simulations (**Fig. 2B**, black dots) as a reference^42^, TG2 binding appeared to produce a reduction in CDR2 exposure in WT models that was less pronounced in R47H models. Together, these results suggest TG2 binding structurally stabilizes the CDR2 loop of WT sTREM2 in a more closed, less solvent-exposed state, while the R47H variant maintains an open, conformationally flexible CDR2 loop that preserves a highly exposed binding interface for TG2.

We next asked whether this more open, binding-accessible CDR2 loop region in R47H sTREM2 yielded significant energetic differences in TG2 binding. To address this, we compared the structural stability, interfacial contact patterns, and per-residue energetics of the WT versus R47H complexes. RMSF analysis of TG2 (**Supplementary Fig. 3B**) revealed that its CH domain remained similarly stable when bound to R47H or WT sTREM2, indicating that the mutation does not induce changes in the conformation of TG2 itself upon binding. In turn, this suggests that TG2-mediated signaling is unlikely to proceed through structural rearrangements within the CH domain but rather through downstream modulation of its C-terminal domain. To assess convergence of the R47H complexes and select representative models for energy analysis, we applied a rolling stability metric identical to that used for WT complexes (see **Methods**; **Supplementary Fig. 3A**). Model 9 showed the highest stability, characterized by interactions between TG2’s PES and a cleft exposed between CDR1 and CDR2 by the open CDR2 loop conformation of R47H sTREM2. Notably, this configuration engages the same general face of sTREM2 as the most stable WT complex, and the open CDR2 loop does not redirect TG2 binding to entirely different surfaces but instead reshapes the local interfacial geometry through exposure of a cleft absent in WT, promoting overall complex stability through enhanced interfacial adaptability.

To determine *how* altered CDR2 accessibility reshaped the interfacial landscape of sTREM2–TG2 binding, we then generated average fractional occupancy profiles for both R47H sTREM2 and TG2. Compared to WT models, TG2 binding to R47H sTREM2 was redistributed: contacts with the Expanded Surface 2 and flexible stalk regions were greatly reduced, while interaction frequencies increased across all CDRs, the hydrophobic tip, and a region spanning residues 90– 120 that does not map to a defined binding site (**Fig. 2C**, left). Diminished TG2 binding to the Expanded Surface 2 in R47H sTREM2 parallels our prior observations on PL binding, reinforcing that this disrupted interaction likely underlies the partial loss-of-function phenotype of the R47H variant unique to sTREM2. Importantly, this observation also extends our previous conclusions beyond PL interactions to include protein receptors. Fractional occupancies of R47H sTREM2 onto TG2 revealed a similar redistribution of contacts across the TG2 surface relative to WT models, most notably decreased contacts with its SMP and increased binding to its PES (**Fig. 2C**, right). Broadly, these changes in interfacial contacts may underlie functional differences in sTREM2–TG2 engagement because of the R47H mutation.

Finally, to directly test whether these structural and interfacial rearrangements translated into energetic differences, we performed per-residue energy decomposition analyses. The results revealed partial overlap in the residues most strongly contributing to the WT and R47H sTREM2– TG2 interfaces: in WT models, R76 provided the dominant favorable contribution, whereas in R47H models this role shifted to the adjacent R77 residue (**Fig. 2D**). This preservation of local arginine-mediated electrostatic anchoring underscores the robustness of the CDR2/Surface 2 motif described for WT complexes. Furthermore, across downsampled trajectories (see **Methods**), no significant difference in the average overall binding free energy was detected between WT and R47H models (**Fig.F**). The most energetically favorable R47H complex (Model 3) featured direct binding between TG2’s PES and sTREM2’s CDR2 loop (**Fig. 2E**), whereas in WT, the lowest-energy conformation engaged primarily through Surface 2 (**Fig. 1C**). Thus, the more closed CDR2 loop conformation observed in WT models does not inherently bind more weakly, but instead favors an alternative TG2 interface, while R47H enhances CDR2 accessibility to PES-mediated binding, yielding comparable overall affinity through a reorganized network of contacts. Notably, comparable binding affinity does not imply equivalent functional engagement — the shift from Surface 2-dominated contacts in WT to CDR2-mediated PES binding in R47H suggests that the functional epitope, rather than binding strength, may be the primary driver of variant-specific differences in TG2 engagement and downstream tau-related outcomes.

### Rational epitope-based optimization of a TG2 therapeutic peptide

Following our analysis of native sTREM2-TG2 interactions, we next sought to translate these residue-level insights into rational peptide design. In their study, Zhang et al. segmented the sTREM2 sequence into four broad fragments and identified residues 77–105 (Peptide 4) as sufficient for TG2 binding by western blotting. They then refined this region by testing shorter sequences within Peptide 4 (Peptides 1-3), ultimately defining Peptide 1 (77–89 aa; RWNGSTAITDDTL), introduced earlier, as the minimal active sequence that best recapitulated full-length TREM2’s protective effects in tauopathy models both *in vitro* and *in vivo* (**Fig. 3A**)^51^.

**Figure 3.**
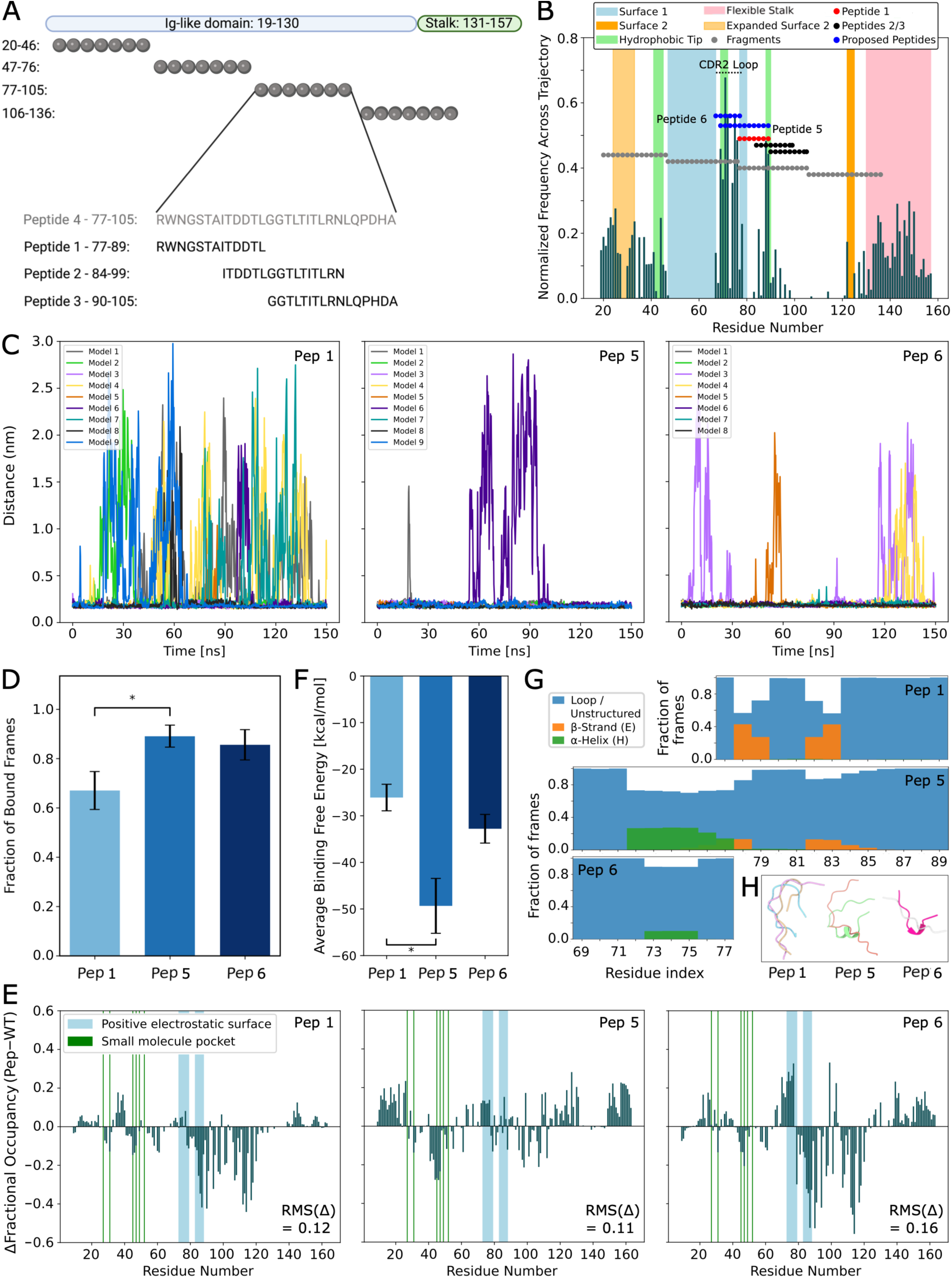
Epitope-guided peptide design stabilizes TG2 binding and recapitulates native sTREM2 interactions. (A) Visualization of the sTREM2-derived peptide design strategy reported by Zhang et al. (B) Normalized fractional occupancy of WT sTREM2 residues by TG2, overlaid with the positions of peptide fragments from Zhang et al. and our proposed peptides. (C) Minimum distance between TG2 and Peptides 1 (left), 5 (center), and 6 (right) plotted over simulation time. (D) Fraction of bound frames for Peptides 1, 5, and 6 over the final 50 ns of the trajectories, defined by a minimum heavy-atom distance ≤ 3.5 Å. Bars show the mean ± SEM; significance was assessed using Welch’s t-tests. (E) Difference in residue-level contact occupancy between each peptide–TG2 complex and the WT sTREM2–TG2 complex, averaged over the final 50 ns of the simulations, with RMS(Δ) values to report the overall deviation from the WT sTREM2 binding profile. (F) MM-PBSA binding free energies for Peptides 1, 5, and 6 bound to TG2, averaged across stably bound models over the final 50 ns of the trajectories. Bars show the mean ± SEM; significance was assessed using Games–Howell multiple-comparison tests. (G) Three-state DSSP secondary structure distributions for Peptides 1 (top), 5 (middle), and 6 (bottom), calculated from TG2-bound simulations and averaged across all models. (H) Representative top conformations identified using the GROMOS clustering method for Peptides 1, 5, and 6 from the TG2-bound simulations.

While Zhang et al.’s optimization protocol was experimentally rigorous, it leaves room for additional mechanistic testing and rational refinement. Building on our earlier observation that Peptide 1 only partially overlaps with sTREM2 regions of highest TG2 occupancy, we next mapped all peptides tested in their study onto the fractional occupancy profiles from our WT sTREM2–TG2 simulations (**Fig. 3B**). As with Peptide 1, Peptides 2 and 3 also fail to fully span both high-occupancy TG2 regions (69–76 aa; 88–89 aa)—though unlike Peptide 1, which narrowly bridges two regions, they only capture singular or partial regions, likely accounting for their weaker activity in the experiments. Consistent with these interaction trends, Peptide 1 emerged as the most effective construct in Zhang et al.’s study, aligning with our WT sTREM2-TG2 interaction profile and further validating the mechanistic basis of our *in silico* approach. This gap in coverage likely reflects a design constraint of their original fragment screen, where the splice site between Peptide 4 and the preceding fragment cuts directly through the highest-occupancy region, preventing its capture in binding assays.

Based on these epitope-mapping insights, we designed two novel sTREM2-derived therapeutic peptide candidates predicted to engage TG2 with improved binding properties: Peptide 5 (69–89 aa; LWLLSFLRRWNGSTAITDDTL) and Peptide 6 (69–77 aa; LWLLSFLRR). Peptide 5 wholly encompasses Peptide 1 while extending N-terminally to include the adjacent high-occupancy segment of the CDR2 loop, whereas Peptide 6 consists solely of the full CDR2-loop sequence (**Fig. 3B**). In addition to the experimentally validated Peptide 1, we performed 150-ns MD simulations of both newly designed peptides in complex with TG2, each initiated from multiple docking conformations (8-9 independent models per peptide, resulting in 26 total simulations). We compared these peptides in terms of their binding stability, TG2 binding-site specificity relative to full-length sTREM2, binding energetics, and structural dynamics. Although we did not assess the therapeutic efficacy of the new designs experimentally, these physics-based simulations provide detailed molecular insight into peptide–TG2 interactions and establish a mechanistic framework for future structure-guided optimization and validation.

To first evaluate the stability of peptide binding to TG2, we calculated the minimum atomic distance between each peptide and TG2 over time (**Fig. 3C**), as well as the RMSD of each peptide relative to TG2 (**Supplementary Figs. 4C, G, K**; see **Supplementary Figs. 4A, E, I** for complex profiles and **Supplementary Figs. 4B, F, J** for TG2 profiles). Together, these metrics quantify whether a peptide remains bound during the simulation and, if so, how much its position deviates from the initial docked structure. This analysis revealed heterogeneous and overall less stable binding behavior for Peptide 1 across simulated models compared to Peptides 5 and 6. In one case (Model 3), Peptide 1 remained stably bound at its initial docking pose, as indicated by consistently low minimum-distance (**Fig. 3C**, left) and RMSD values (**Supplementary Fig. 4C**) throughout the simulation. In several other Models (2, 5, 6, 8, and 9), Peptide 1 transiently dissociated but subsequently re-stabilized at an alternative TG2 site, reflected by brief excursions to high minimum distances followed by elevated yet steady RMSD values upon re-binding (**Fig. 3C**, left; **Supplementary Fig. 4C**). In the remaining Models (1, 4, and 7), Peptide 1 completely dissociated and failed to stably rebind, characterized by persistently high or fluctuating minimum distances and unstable RMSD traces, particularly during the final 50 ns. In contrast, both Peptide 5 and Peptide 6 exhibited generally lower RMSD values (**Supplementary Figs. 4G,** K) and far fewer excursions away from the TG2 surface, indicating improved binding stability (**Fig. 3C**, middle for Peptide 5 (Models 1, 6); right for Peptide 6 (Models 3-5)). While both *in silico*-derived peptides underwent occasional binding and re-binding events across trajectories, Peptide 5 remained bound in all nine models during the final 50 ns, and Peptide 6 dissociated in only two of eight models during this period (Models 3, 4).

To more rigorously compare binding stability, we next quantified peptide residence fraction, or the fraction of trajectory frames (averaged across all models over the final 50 ns) in which each peptide remained bound to TG2, defined as maintaining a minimum distance within 3.5 Å (**Fig. 3D**). To ensure this metric reflected sustained rather than transient interactions, only contacts persisting for at least 5 ns were included. Consistent with the RMSD and minimum-distance analyses, Peptide 5 remained bound for a significantly greater fraction of time than Peptide 1 (t = −2.478; p = 0.028; n = 9). While Peptide 6 also exhibited a higher mean bound fraction than Peptide 1, this difference was not statistically significant. One explanation for these differences in residency might be the length of the peptides. Longer peptides, like Peptide 5 (21 aa), contain more residues that can create stable contacts with TG2, compared to shorter Peptides 1 and 6 (13 and 9 aa, respectively). However, therapeutic peptides are generally under 40 amino acids in length, and all three of these therapeutic candidates meet this requirement. Furthermore, these results indicate that while the originally reported Peptide 1 can interact with TG2, its binding is relatively weak and unstable; in contrast, our rationally designed peptides—particularly Peptide 5—demonstrated markedly enhanced binding persistence across independent simulations.

While identifying peptides that bind TG2 stably is essential, stability alone is insufficient to ensure therapeutic relevance if binding occurs at off-target or nonfunctional regions. To this end, we posited that peptides most closely recapitulating the binding profile of full-length sTREM2 (**Fig. 1F**, right) would best mimic its neuroprotective interactions with TG2 and thus represent stronger therapeutic candidates. Notably, although TG2 lacks a clearly defined catalytic or receptor-binding pocket, our earlier results—together with findings from Wu et al.—highlight conserved surface regions (SMP and PES) that are likely to mediate downstream signaling. To assess site specificity, we calculated residue-level contact frequencies between each peptide and TG2 (averaged across all models over the final 50 ns; **Supplementary Fig. 5A**) relative to those observed in full-length WT sTREM2–TG2 simulations (Δoccupancy; **Fig. 3E**). Qualitatively, all three peptides showed mild reductions in contact frequency at the SMP. More notably, Peptide 1 exhibited a pronounced loss of binding at the PES (observed by the negative occupancies in **Fig. 3E**, left). In contrast, Peptide 5 more effectively preserved native PES contacts, whereas Peptide 6 displayed a mixed pattern of increased contacts with some PES residues and decreased contacts with others (**Fig. 3E**, middle and right, resp.). Peptide 5 also displayed increased interactions with TG2’s unstructured termini. However, the biological relevance of this observation remains uncertain, as these regions lack stabilizing contacts in the crystal structure and thus remained flexible in the simulations, while in a membrane-associated context TG2’s termini would likely be more conformationally restrained.

To collectively quantify how these alterations in peptide–TG2 interactions affect overall similarity to full-length WT sTREM2’s binding profile, we calculated the root-mean-square deviation of each peptide’s per-residue occupancy values relative to those of WT sTREM2 (RMS(Δ); **Fig. 3E**). Peptide 5 exhibited the lowest RMS(Δ) (0.11), indicating that its binding profile most closely recapitulates native sTREM2 contacts, trailed by Peptide 1 (0.12) and Peptide 6 (0.16). Although peptide length may partially influence this metric—as Peptide 5 is the longest, followed by Peptide 1 and then Peptide 6—the fact that Peptide 5 selectively preserves PES interactions rather than uniformly increasing contacts across all TG2 regions suggests that its improved mimicry of WT sTREM2 arises from targeted engagement rather than increased length alone.

We next asked whether the observed differences in peptide binding stability and TG2 binding-site specificity—both relative to WT sTREM2 and between the peptides themselves—translated into meaningful distinctions in binding energetics. To test this, we performed MM-PBSA calculations on the peptide–TG2 complexes over the final 50 ns of each simulation, restricting the analysis to models in which the peptides remained stably bound during this period and averaging results across these trajectories. Peptide 5 exhibited the most favorable mean binding free energy, followed by Peptide 6 and Peptide 1 (**Fig. 3F**). Because the groups had unequal sample sizes and substantially different variances, pairwise significance was assessed using Games–Howell multiple-comparison tests. Peptide 5 exhibited significantly more favorable binding than Peptide 1 (p = 0.011; n = 9 and 6, respectively), whereas differences between Peptide 1 and Peptide 6 and between Peptide 5 and Peptide 6 were not statistically significant. Together, these energetic trends indicate that both rationally designed peptides favor more favorable TG2 binding than Peptide 1, with the significant Peptide 5–Peptide 1 difference providing the strongest evidence for enhanced relative binding energetics in the Peptide 5 design. Perhaps it is the combination of these three factors that can enhance sTREM2-dependent TG2 signaling in ways that mitigate disease pathology. The complexity of this interaction, and the therapeutic benefits of targeting specific residues, remains an active area of investigation that can be further elucidated with our *in silico* peptides.

Finally, we examined whether structural features intrinsic to each peptide might contribute to their observed differences in binding stability, energetics, and specificity. Using dictionary of secondary structure in proteins (DSSP) classification, we quantified the secondary-structure dynamics of each peptide across both unbound trajectories (**Supplementary Fig. 5B**) and TG2-bound complexes (**Fig. 3G**), calculating the fraction of total simulation time during which each residue adopted unstructured loop/turn, β-strand, or α-helical conformations. As expected for short, flexible peptides, all three exhibited similar, largely disordered conformations with minimal persistent β-strand or α-helical content (<40% of total simulation time for either ordered element). Secondary-structure profiles were also similar between bound and unbound states, indicating that peptide conformation was not strongly induced or altered upon TG2 binding within the simulated timescales. Across overlapping sequence regions, the peptides showed similar overall secondary-structure propensities, with only minor differences in the extent and persistence of each element. Peptides 1 and 5 both exhibited low-persistence β-strand character between residues 78–84, though this feature was slightly more stable in Peptide 1. Peptides 5 and 6 also formed α-helical structure at their newly added, overlapping N-terminal residues (73–76 aa), with the helix in Peptide 5 extending slightly further and persisting marginally longer than in Peptide 6. Although these differences were modest, the shared helical propensity of Peptides 5 and 6 is notable because these residues correspond to the CDR2 loop of full-length sTREM2, where the same region adopts partial α-helical structure that is further stabilized by the R47H AD-risk mutation. This suggests that the additional N-terminal residues in Peptides 5 and 6 may better preserve the native conformational preferences of the CDR2 region, making them more structurally faithful mimetics of the parent sTREM2 epitope than Peptide 1.

These structural trends are also visually apparent in the top clustered conformations for each peptide (**Fig. 3H**). Partial preorganization of the CDR2-derived segment may favor TG2 engagement by reducing the conformational rearrangement required for binding and stabilizing interfacial residue orientation. Although no secondary-structure restraints were imposed, the observed helical propensity may therefore contribute to peptide–TG2 binding. However, because these differences were modest and similar in bound and unbound states, the simulations do not establish a causal relationship between helix formation and enhanced binding. Likewise, the transient secondary structure does not by itself indicate increased self-association or fibrillization propensity.

Taken together, these analyses show that rational epitope-based design improved multiple features of TG2 engagement, including binding persistence, native-like contact patterns, and binding energetics. Peptide 5 emerged as the strongest candidate across these metrics, supporting the value of incorporating the adjacent high-occupancy CDR2 segment. Beyond the candidates themselves, these peptides function as a reductionist probe of the native sTREM2–TG2 interaction: by isolating discrete epitope segments, they indicate that the CDR2 region contributes disproportionately to productive TG2 engagement relative to its length, and that PES contacts are better preserved when that segment is included.

## Conclusions

In summary, our simulations define a conserved, electrostatically driven interface underlying the sTREM2–TG2 interaction, centered on a novel positive electrostatic surface (PES) on TG2 and a CDR2-based binding motif in sTREM2 which consists of charged anchoring residues with flanked by a region of enriched hydrophobic content. We show that the AD-risk R47H variant preserves overall binding affinity while reorganizing interfacial contacts toward the PES through maintenance of an open, solvent-exposed CDR2 loop. Because R47H’s established loss-of-function is directed at membrane-bound TREM2, a comparatively spared sTREM2–TG2 axis offers one explanation for reports that R47H does not exacerbate, and may attenuate, tau pathology in tauopathy-only models. Guided by these residue-level insights, we propose novel TG2-targeting therapeutic peptide candidates and demonstrate that epitope-based optimization increases the fraction of simulation time spent bound to TG2, yields more favorable binding energetics, and more closely recapitulates native sTREM2–TG2 contact patterns than the previously reported sTREM2(77–89) peptide. Uniquely, our approach defines peptide boundaries directly from residue-level occupancy maps of the native complex, capturing a high-occupancy region that the splice site of the original fragment screen bisected, and engineers candidates against a dynamic conformational ensemble rather than a single static structure. Together, these results establish a mechanistic framework for the sTREM2–TG2 axis and provide a structure-guided foundation for future optimization and experimental validation of TG2-targeting peptides.

Several limitations should be noted. Our conclusions are based on a rigorous but fully computational framework, and while we consistently relate our findings to prior experimental observations, direct experimental testing will be essential to validate the proposed mechanisms and peptide designs. In addition, our models use a truncated TG2 construct, omit N-linked glycosylation at N20 and N79, the latter near CDR2, which in endogenous sTREM2 could affect TG2 accessibility, rely on finite-timescale MD simulations and MM-PBSA energetics, and treat peptides as free in solution. Each peptide was also simulated as a single copy, so assessing self-association would require multi-copy simulations or sequence-based aggregation prediction. Incorporating membrane context, enhanced sampling strategies, and more rigorous free-energy methods will therefore be important in future work. Moreover, information on TG2 expression and regulation in neurons remains limited. While Zhang et al. demonstrated a critical role for TG2 in mediating sTREM2’s effects on tau pathology, substantial opportunities remain to investigate how TG2 expression and role across neuronal subtypes and disease stages. Finally, peptide structural behavior may also be influenced by delivery strategy: in the study by Zhang et al., the peptide was fused to a modified HIV Tat sequence to enable blood–brain barrier permeability, and such tags may affect peptide folding and TG2 accessibility.

Beyond the specific candidates proposed here, our work demonstrates how residue-level interaction maps derived from full-length protein–protein simulations can define peptide boundaries that fragment-based screening may systematically overlook. By treating the native complex as the design template, this approach engineers candidates against dynamic conformational ensembles rather than a single static structure, and helps identify the sequence and structural features that likely underlie productive TG2 engagement. More broadly, our findings position the sTREM2–TG2 axis as a mechanistically tractable node linking microglial and neuronal contributions to tau pathology. Therapeutically, this axis engages distinct biochemical and pathological features from strategies targeting membrane-bound TREM2.

Whether epitope-selective engagement translates into differential TG2 activation, and whether the interfacial rearrangement we observe in R47H carries functional consequences, remain active areas of investigation that our *in silico* framework is well positioned to guide.

## Acknowledgements

This work utilized the Alpine high-performance computing resource and the Blanca condo computing resource at the University of Colorado Boulder. Alpine is jointly funded by the University of Colorado Boulder, the University of Colorado Anschutz, and Colorado State University. Blanca is jointly funded by computing users and the University of Colorado Boulder. This material is based upon work supported by the National Science Foundation Graduate Research Fellowship Program under Fellow ID (2024361899). Any opinions, findings, conclusions, or recommendations expressed in this material are those of the author(s) and do not necessarily reflect the views of the National Science Foundation.

## Use of AI Technology

During the preparation of this manuscript, the authors used Claude (Anthropic) and ChatGPT (OpenAI) to assist with editing and refining the wording of the manuscript. No data, figures, or scientific interpretations were generated by these tools. The authors reviewed and edited all text, and take full responsibility for the accuracy and integrity of the content of the publication.

## Data Availability

The current manuscript is a computational study, so no experimental data have been generated for this manuscript. All analyzed computational data is included in the main figures and supplementary figures. Processed trajectories for production simulations and related structure files are available upon reasonable request from the corresponding author.

