## Supplementary Material for "Investigating the therapeutic potential of soluble TREM2 interactions with transgelin-2 in Alzheimer’s disease"

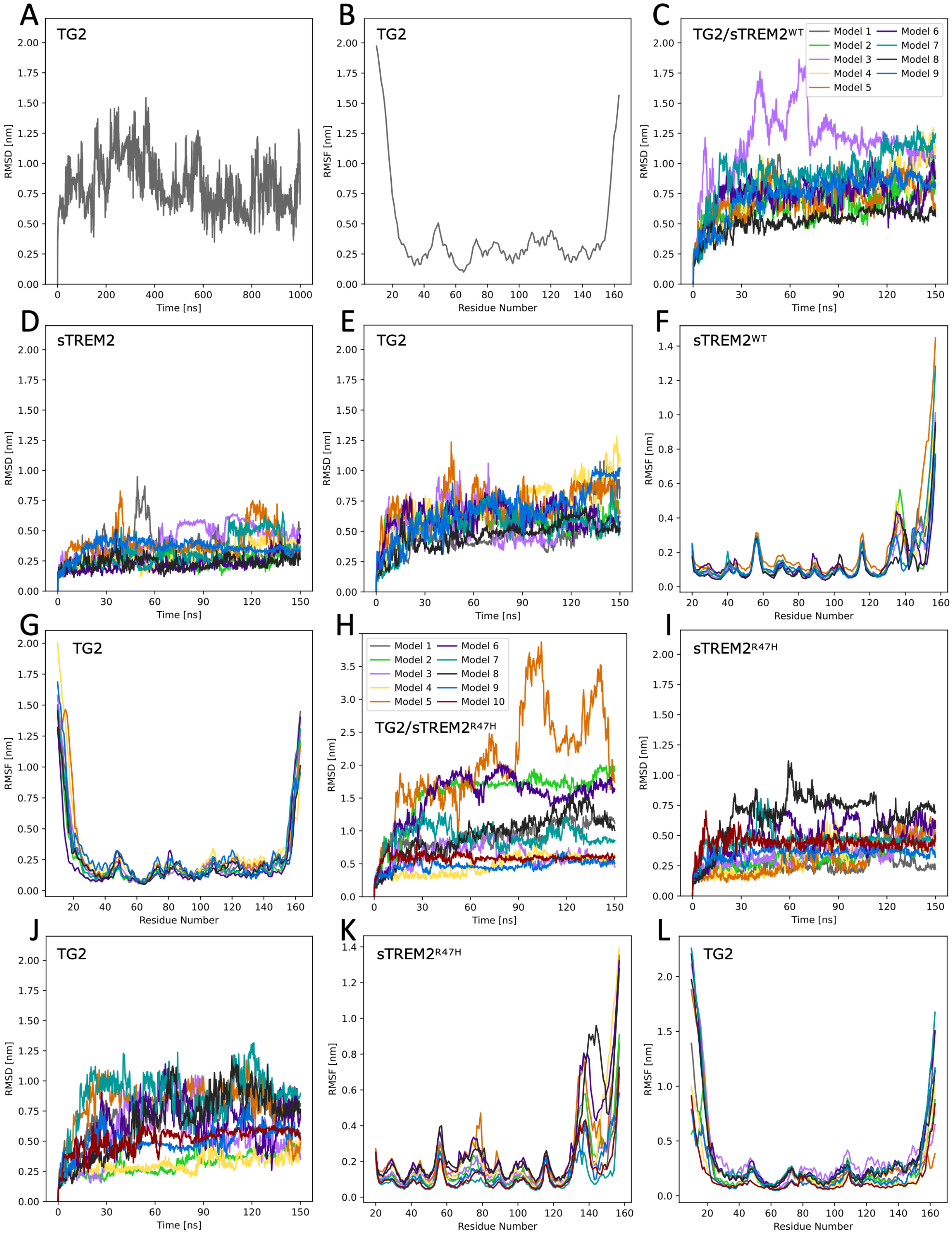


**Figure S1. Comparative Cα-RMSD and Cα-RMSF analyses across individual sTREM2–TG2 MD simulations.** (A) TG2 RMSD in solvent. (B) TG2 RMSF in solvent. (C) Whole-system

RMSD of TG2/WT sTREM2 complexes. (D) sTREM2 RMSD in TG2/WT sTREM2 complexes. (E) TG2 RMSD in TG2/WT sTREM2 complexes. (F) sTREM2 RMSF in TG2/WT sTREM2 complexes. (G) TG2 RMSF in TG2/WT sTREM2 complexes. (H) Whole-system RMSD of TG2/R47H sTREM2 complexes. (I) sTREM2 RMSD in TG2/R47H sTREM2 complexes. (J) TG2 RMSD in TG2/R47H sTREM2 complexes. (K) sTREM2 RMSF in TG2/R47H sTREM2 complexes. (L) TG2 RMSF in TG2/R47H sTREM2 complexes.

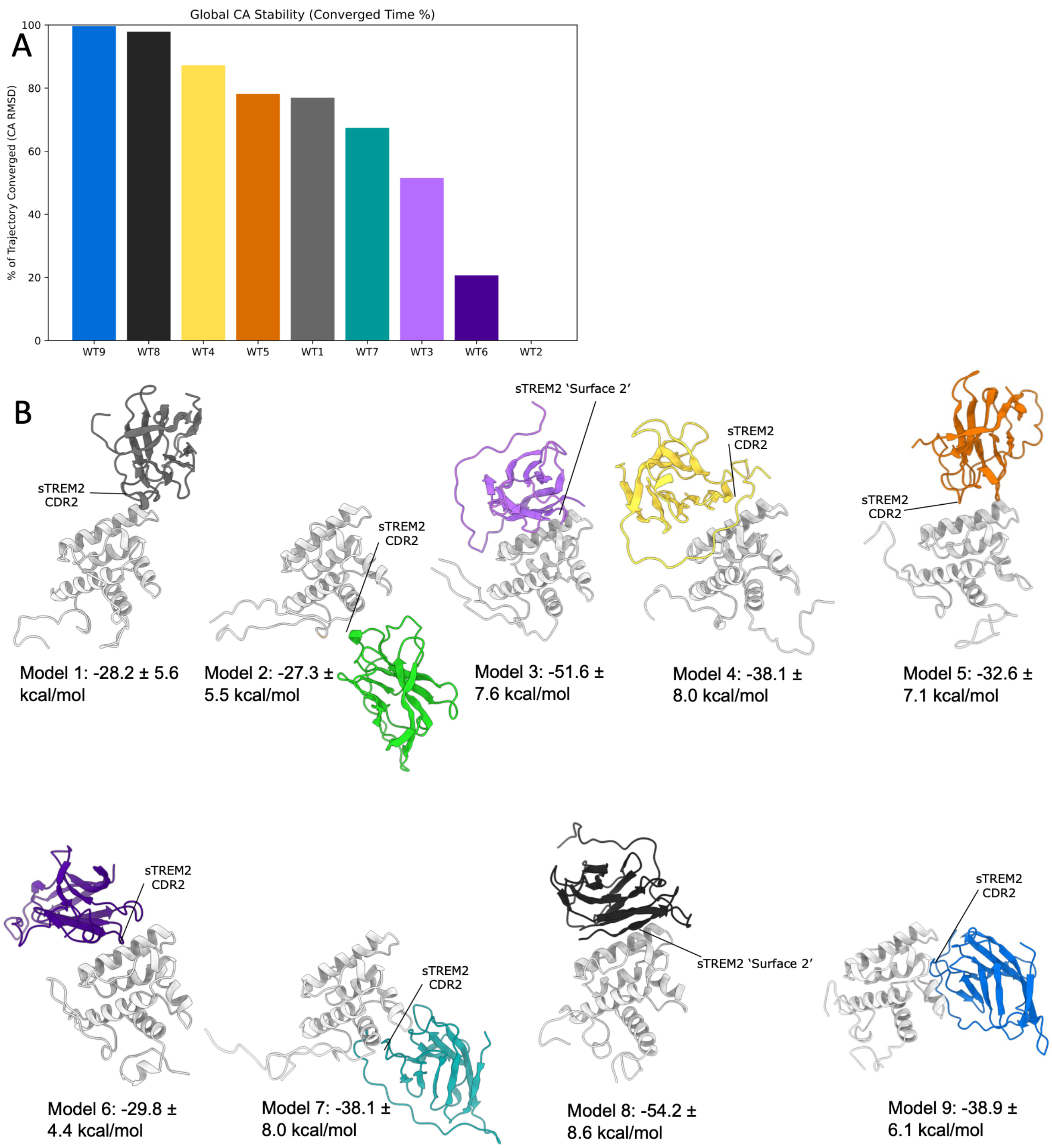


**Figure S2. Stability and energetics of WT sTREM2–TG2 models.** (A) Global Cα stability of WT sTREM2–TG2 complexes, quantified as the percentage of simulation time each trajectory remained within a ±3 Å Cα RMSD window relative to the initial docked pose (see Methods), assessed in 20 ns intervals. (B) Representative structures from the final 50 ns of each WT sTREM2-TG2 simulation, with TG2 shown in light gray and sTREM2 colored by model. The sTREM2 CDR2 and alternative 'Surface 2' binding modes are indicated. Reported values are MMPBSA binding scores (mean ± SD, kcal/mol).

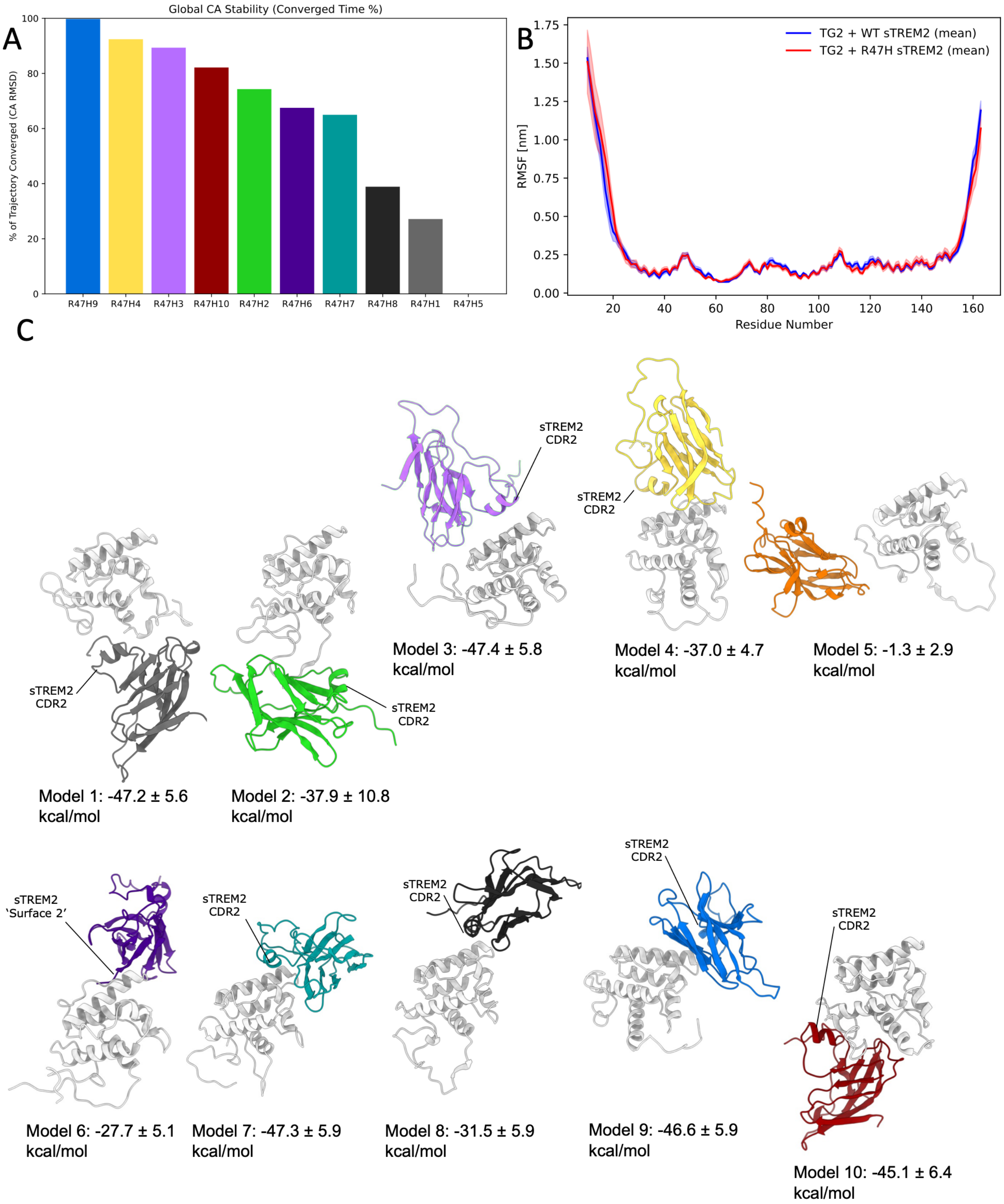


**Figure S3. Stability and energetics of R47H sTREM2–TG2 models.** (A) Global Cα stability of R47H sTREM2–TG2 complexes, quantified as the percentage of simulation time each trajectory remained within a ±3 Å Cα RMSD window relative to the initial docked pose (see Methods), assessed in 20 ns intervals. (B) Cα RMSF profiles for TG2 bound to WT (blue) and R47H (red) sTREM2, shown as the mean ± SEM across independent simulations. (C) Representative structures from the final 50 ns of each R47H sTREM2-TG2 simulation, with TG2 shown in light gray and sTREM2 colored by model. The sTREM2 CDR2 and alternative ‘Surface 2’ binding modes are indicated where possible. Reported values are MMPBSA binding scores (mean ± SD, kcal/mol).

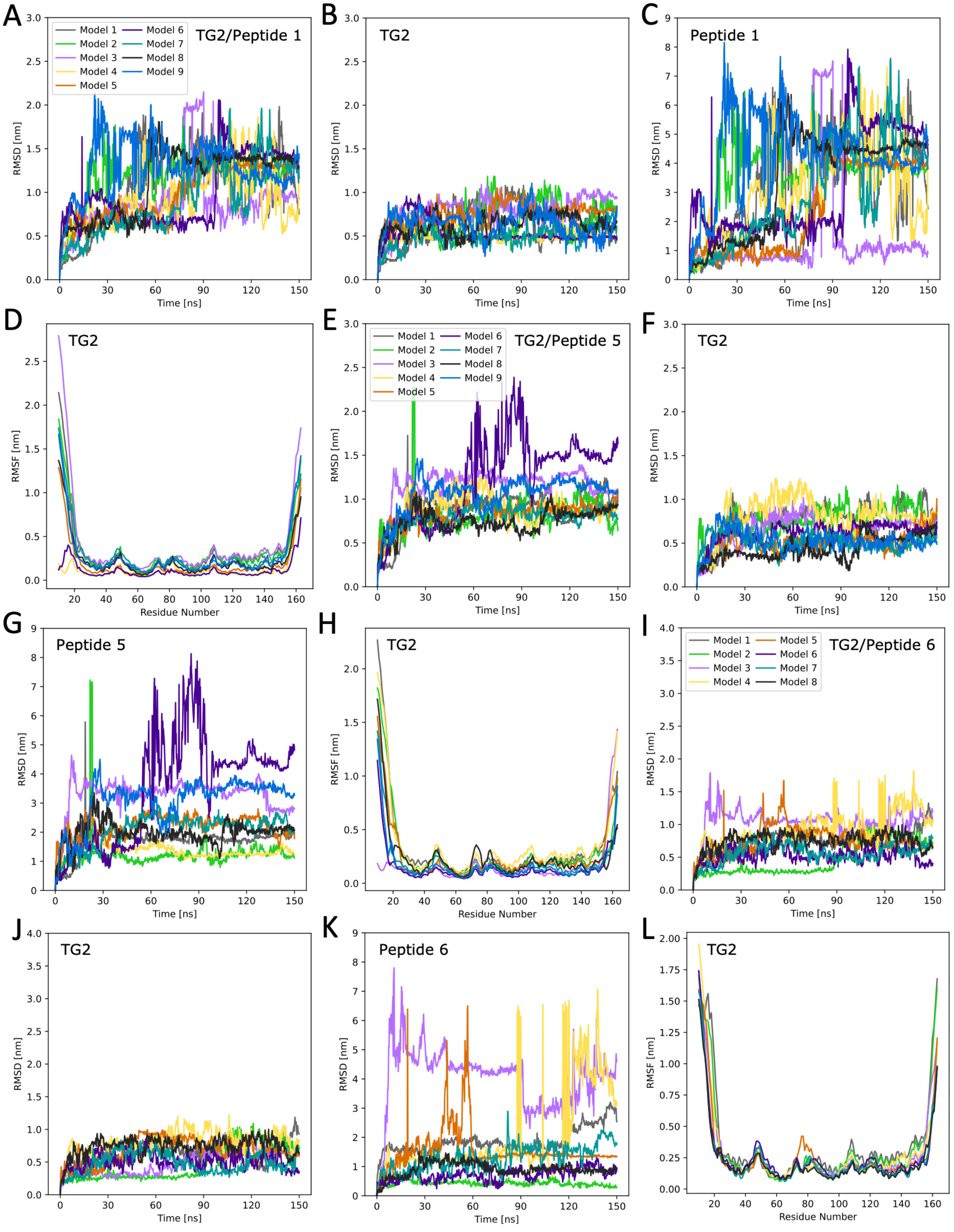


**Figure S4. Comparative Cα-RMSD and Cα-RMSF analyses across individual TG2–peptide MD simulations.** (A) Whole-system RMSD of TG2/Peptide 1 complexes. (B) TG2 RMSD in TG2/Peptide 1 complexes. (C) Peptide 1 RMSD in TG2/Peptide 1 complexes, calculated relative to TG2. (D) TG2 RMSF in TG2/Peptide 1 complexes. (E) Whole-system RMSD of TG2/Peptide

5 complexes. (F) TG2 RMSD in TG2/Peptide 5 complexes. (G) Peptide 5 RMSD in TG2/Peptide 5 complexes, calculated relative to TG2. (H) TG2 RMSF in TG2/Peptide 5 complexes. (I) Whole-system RMSD of TG2/Peptide 6 complexes. (J) TG2 RMSD in TG2/Peptide 6 complexes. (K) Peptide 6 RMSD in TG2/Peptide 6 complexes, calculated relative to TG2. (L) TG2 RMSF in TG2/Peptide 6 complexes.

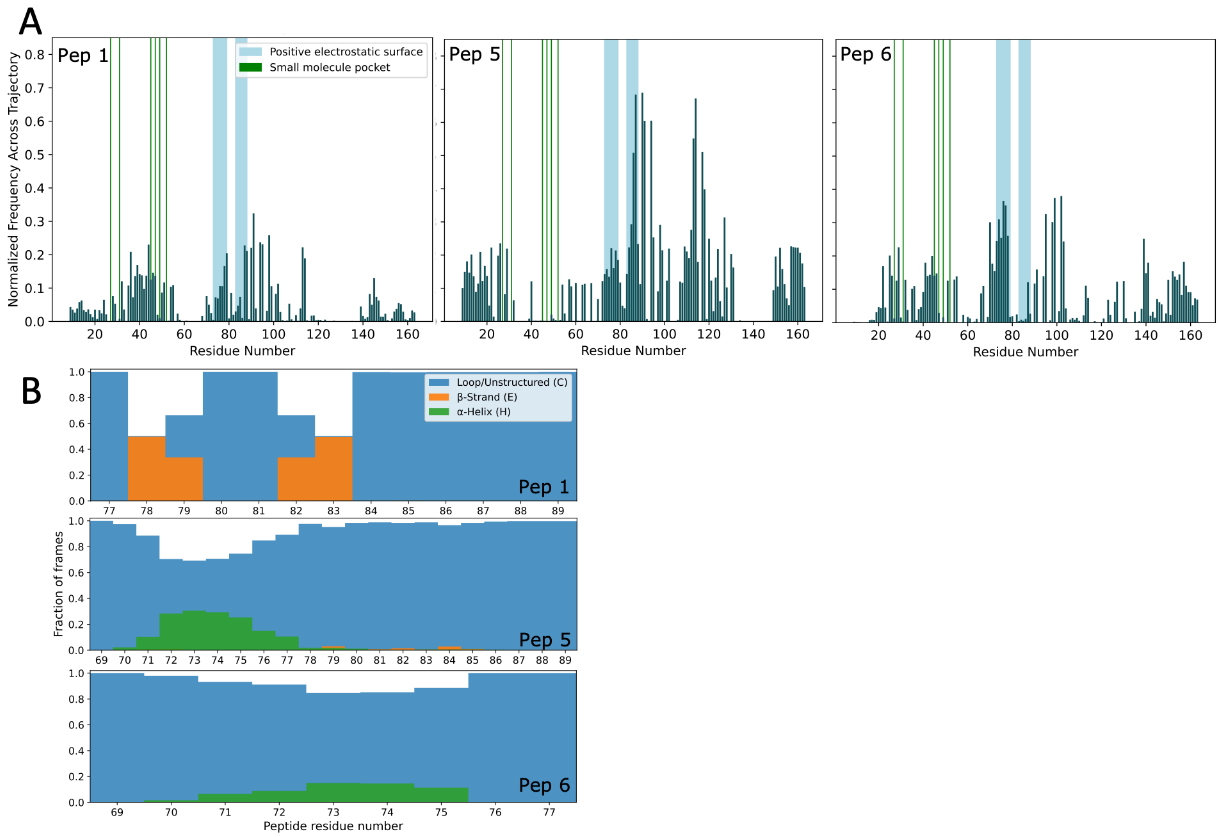


**Figure S5. Peptide fractional occupancy and secondary structure analysis.** (A) Normalized fractional occupancy of TG2 residues by Peptides 1 (left), 5 (middle), and 6 (right). (B) Three-state DSSP secondary structure distributions for Peptides 1 (top), 5 (middle), and 6 (bottom), calculated from 1 μs simulations of each peptide in solvent.
